# Differential requirement of NPHP1 for compartmentalized protein localization during photoreceptor outer segment development and maintenance

**DOI:** 10.1101/2021.01.20.427412

**Authors:** Poppy Datta, J. Thomas Cribbs, Seongjin Seo

## Abstract

Nephrocystin (NPHP1) is a ciliary transition zone protein and its ablation causes nephronophthisis (NPHP) with partially penetrant retinal dystrophy. However, the precise requirements of NPHP1 in photoreceptors are not well understood. Here, we characterize retinal degeneration in a mouse model of NPHP1 and show that NPHP1 is required to prevent infiltration of inner segment plasma membrane proteins into the outer segment during the photoreceptor maturation. We demonstrate that *Nphp1* gene-trap mutant mice, which were previously described as null, are in fact hypomorphs due to the production of a small quantity of functional mRNAs derived from nonsense-associated altered splicing and skipping of two exons including the one harboring the gene-trap. In homozygous mutant animals, inner segment plasma membrane proteins such as syntaxin-3 (STX3), synaptosomal-associated protein 25 (SNAP25), and interphotoreceptor matrix proteoglycan 2 (IMPG2) accumulate in the outer segment when outer segments are actively elongating. This phenotype, however, is spontaneously ameliorated after the outer segment elongation is completed. Retinal degeneration also occurs temporarily during the photoreceptor maturation but stops afterward. We further show that *Nphp1* genetically interacts with *Cep290*, another NPHP gene, and that a reduction of *Cep290* gene dose results in retinal degeneration that continues until adulthood in *Nphp1* mutant mice. These findings demonstrate that NPHP1 is required for the confinement of inner segment plasma membrane proteins during the outer segment development, but its requirement diminishes as photoreceptors mature. Our study also suggests that additional mutations in other NPHP genes may influence the penetrance of retinopathy in human NPHP1 patients.

## Introduction

Photoreceptor cells in the retina are structurally and functionally compartmentalized. Proteins that transduce light into chemical and electrical signals are confined to an apical compartment called the outer segment (OS). This structure is an elaborate modification of primary cilia, which exist in most cells in vertebrates (see [1, 2] for a comprehensive review). In contrast, energy production and protein/lipid synthesis occur in another compartment, the inner segment (IS). Constituents of the OS are synthesized in the IS and transported to the OS through a narrow channel, the connecting cilium. The connecting cilium is equivalent to the transition zone in primary cilia and plays critical roles in regulating protein trafficking and confinement between the IS and the OS [3].

Nephrocystin (also known as NPHP1) is a protein that localizes to the connecting cilium in photoreceptors and the transition zone in primary cilia [4-9]. At the transition zone, NPHP1 is a part of a multi-protein complex that functions as a gate to control protein trafficking in and out of the ciliary compartment [8-13]. Ablation of individual components in this complex causes a group of related diseases, including nephronophthisis (NPHP), Joubert syndrome (JBTS), and Meckel-Gruber syndrome (MKS) [14, 15]. Retinal anomalies are common in patients with these diseases. While *NPHP1* is primarily associated with NPHP [16-19], a subset (6-10%) of NPHP1 patients exhibits retinal dystrophy as an extra-renal manifestation [19, 20]. This suggests a role of NPHP1 in photoreceptors. However, its precise functions in photoreceptors and the molecular/genetic basis of the incomplete penetrance are not well understood.

Two NPHP1 mouse models have been described thus far. In one model (*Nphp1*^*del20*^), the last exon (exon 20) was deleted, and homozygous mutant animals displayed severe retinal degeneration with OS morphogenesis defects [5, 21]. *Nphp1*^*del20/del20*^ animals also exhibited male infertility due to a defect in spermatogenesis [21]. The second model (*Nphp1*^*neo*^; hereafter referred to as *Nphp1*^*gt*^) was generated by an insertion of a gene-trap (a *PGK-neo-pA* cassette) into exon 4 (**S1 Fig**) [22]. This mutation is expected to generate a null allele because it disrupts the reading frame of *Nphp1*. However, retinal degeneration in this model is significantly milder than that of *Nphp1*^*del20*^ mice: a slight reduction of the photoreceptor cell layer was observed at postnatal (P) day P21 and 2 months of age, and rhodopsin (RHO) trafficking was only marginally affected. Retinal phenotypes in older animals were not reported. The basis of this phenotypic difference in severity between these two mouse models is unknown, but disparities in the genetic background were suggested as a potential contributing factor [22]. Currently, only the *Nphp1*^*gt*^ model is available to the research community and has been used in several studies to investigate genetic interactions with other ciliopathy genes [9, 23, 24].

We previously proposed that the OS acts as a sink for membrane proteins because of its large size, high membrane content, and continuous renewal [3]. This also suggests that accumulation of IS membrane proteins in the OS would be a common pathomechanism of retinal degenerations associated with ciliary gate defects. Indeed, IS plasma membrane-associated proteins including syntaxin-3 (STX3), syntaxin-binding protein 1 (STXBP1), synaptosomal-associated protein 25 (SNAP25), and interphotoreceptor matrix proteoglycan 2 (IMPG2) rapidly accumulate in the OS when the function of CEP290, another component of the ciliary gate complex, is compromised [25]. Notably, inactivating mutations in *CEP290* cause Leber congenital amaurosis (LCA), an early-onset severe retinal degeneration, in humans [26-29]. In this work, we sought to determine the requirement of NPHP1 for protein confinement in photoreceptors and test whether the aforementioned pathomechanism underlies retinal degeneration in *NPHP1*-associated retinopathies.

## Materials and methods

### Mouse and genotyping

*Nphp1*^*gt*^ and *Cep290*^*fl*^ mice were previously described [22, 30] and obtained from the Jackson laboratory (*Nphp1*^*tm1Jgg*^*/J*, #013169; *Cep290*^*tm1Jgg*^*/J*, #013701). *iCre75* mice were a generous gift from Dr. Ching-Kang Chen [31]. The lack of *rd1* mutation (in *Pde6b*) was confirmed by PCR using primers described in [25]. The *rd8* mutation (in *Crb1*) was eliminated by breeding. All animals used in this study were *Pde6b*^*+/+*^*;Crb1*^*+/+*^. For genotyping, mouse tail snips were collected at the time of weaning (P19-P24) or after euthanasia. Tail DNAs were extracted by Proteinase K digestion (Sigma-Aldrich; RPROTK-RO) in Tail Lysis Buffer (10 mM Tris pH 8.0, 100 mM NaCl, 10 mM EDTA, 1% SDS, 0.3 mg/ml Proteinase K) followed by ethanol precipitation. Genotyping was conducted by PCR using GoTaq G2 Flexi DNA polymerase (Promega) and primers listed in **Table 1**. PCR protocols are available upon request. All animals were maintained in 12-hour light/dark cycles and fed *ad libitum* standard mouse chow. All animal procedures were approved by the Institutional Animal Care and Use Committee (IACUC) of the University of Iowa (Protocol#: 8011301) and conducted following the recommendations in the Guide for the Care and Use of Laboratory Animals of the National Institutes of Health.

**Table 1.**
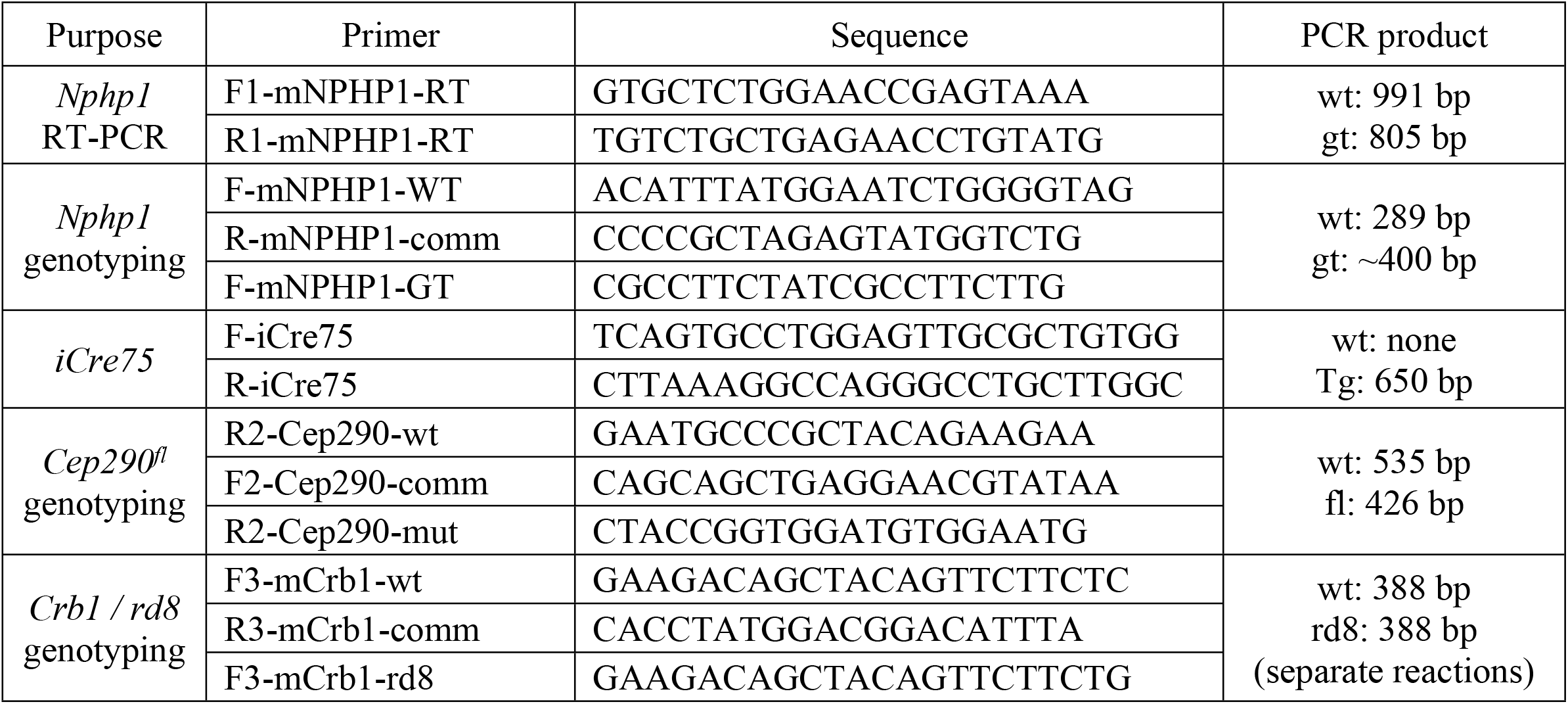
Primers used in this study.

### Plasmids and antibodies

A full-length mouse *Nphp1* coding sequence (BC118953) was obtained from Transomic Technologies. FLAG-tagged NPHP1 expression vectors (pCS2FLAG-NPHP1 (full-length), pCS2FLAG-NPHP1 Δ(49-110), and pCS2FLAG-NPHP1 aa242-685) were generated by inserting PCR-amplified DNA fragments into the EcoRI-XhoI sites of the pCS2FLAG plasmid, which was produced by replacing the 6x Myc tag with a 3x FLAG tag in the pCS2+MT vector [32], using a GenBuilder cloning kit (GenScript). pCS2HA-NPHP1 and pCS2HA-NPHP1 Δ(49-110) were generated by inserting the same PCR fragments into the pCS2HA vector [25]. Expression vectors pEGFP-NPHP2 and pEGFP-NPHP5 were generous gifts from Drs. Val C. Sheffield and William Y. Tsang, respectively. All constructs were sequence-verified by Sanger sequencing. Antibodies used for immunoblotting and immunohistochemistry are described in **Table 2**.

**Table 2.**
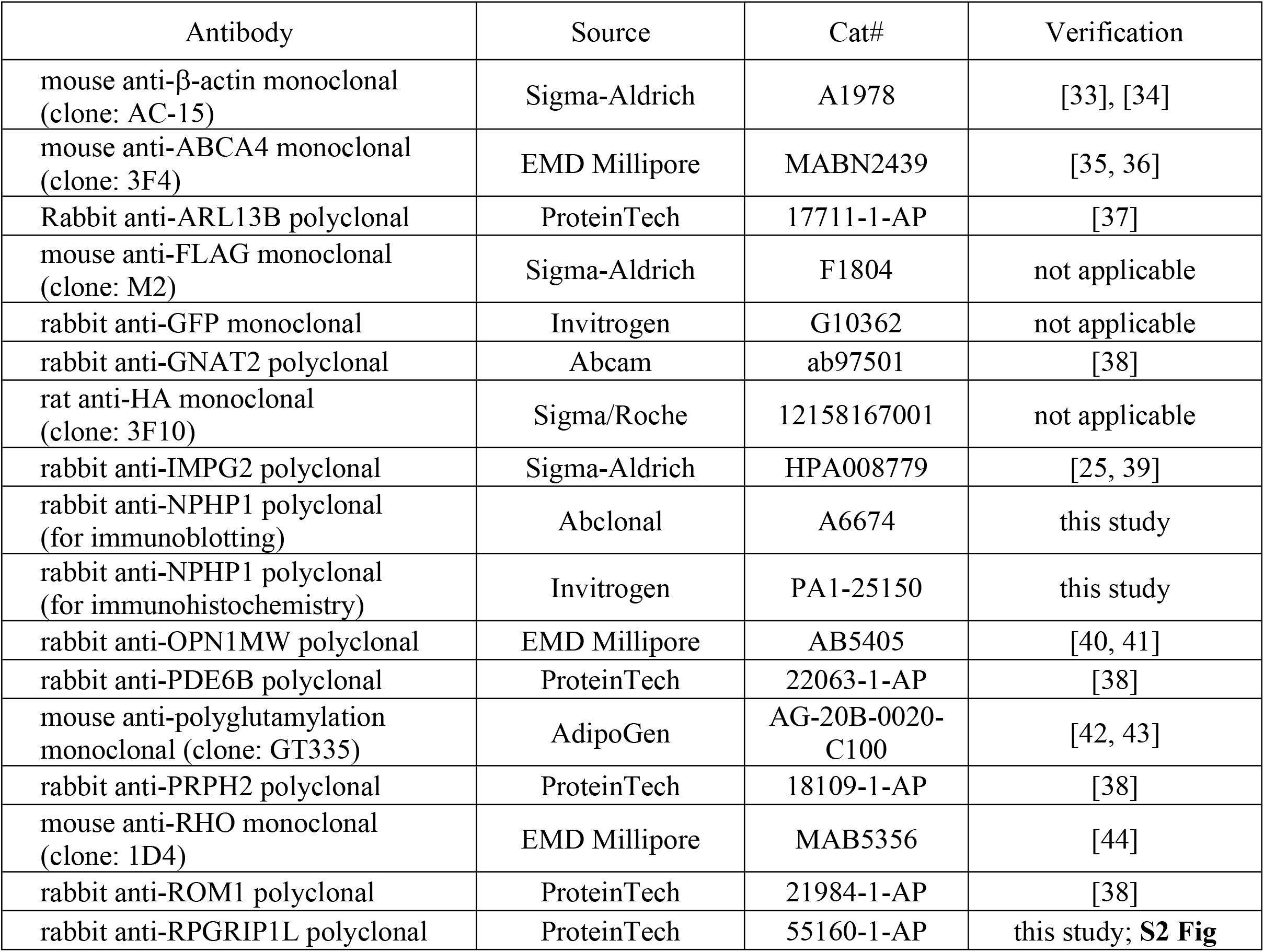
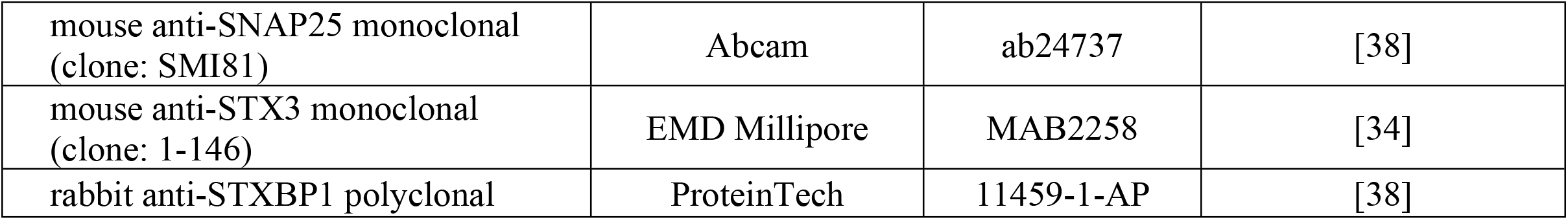
Antibodies used in this study.

### Immunoblotting

To extract proteins from mouse testes, animals were euthanized by CO_2_ asphyxiation followed by cervical dislocation, and testicles were removed. Collected testicles were homogenized with Polytron PT 1200E (Kinematica) in an ice-cold lysis buffer (50 mM HEPES pH7.0, 150 mM NaCl, 2 mM MgCl_2_, 2 mM EGTA, 1% Triton X-100) supplemented with Protease Inhibitor Cocktail (Bimake). Homogenates were clarified by centrifugation at 20,000x g for 15 min at 4 °C and supernatants were collected. Protein concentrations were measured using a DC Protein Assay kit (Bio-Rad) with BSA protein standards and AccuSkan GO spectrophotometer (Fisher Scientific). Fifty μg of proteins were mixed with NuPAGE LDS Sample Buffer (Invitrogen) and Reducing Agent (Invitrogen) and loaded on a 4-12% (wt/vol) NuPAGE Bis-Tris gel (Invitrogen). After electrophoresis, proteins were transferred to a 0.45-μm nitrocellulose membrane (Bio-Rad). Immunoblotting was performed following standard protocols, and primary antibodies used are listed in **Table 2**. Proteins were detected with horseradish peroxidase (HRP)-linked anti-mouse IgG and anti-rabbit IgG secondary antibodies (Cell Signaling) and SuperSignal West Dura Extended Duration Substrate (Thermo Scientific). Images were taken with a ChemiDoc system (Bio-Rad).

### Immunohistochemistry

Mouse eyes were collected, fixed, and embedded in Neg-50 Frozen Section Medium (Richard-Allan Scientific) as previously described [25]. Frozen eyecups were mounted on a cryostat chuck in an orientation that sections became perpendicular to the retinal plane at the central retina. Eight-μm thick, serial sections were collected from the middle 1/3 of eyecups and used for immunohistochemistry. Immunostaining and imaging procedures were previously described elsewhere [25]. To assess the photoreceptor cell loss, 3 serial sections that contained the central retina were selected and the number of rows of photoreceptor cell nuclei was counted near the center of the retina (300-600 μm from the optic nerve head; 2 locations/section). One-way ANOVA followed by Tukey’s multiple comparison test was used for statistical analyses using GraphPad Prism software. *P* values smaller than 0.01 were regarded as statistically significant.

Retinal sections to assess NPHP1 localization to the connecting cilium were prepared as previously described [25]. Fluorescence intensities were measured using ImageJ/Fiji. Background fluorescence intensity in the vicinity of each connecting cilium was subtracted from the intensity of NPHP1 at the connecting cilium. Two-tailed *t*-test assuming unequal variances was performed for statistical analysis.

mIMCD-3 cells were obtained from ATCC (#CRL-2123) and cultured in DMEM:F12 Medium (Invitrogen) supplemented with 10% fetal bovine serum (Sigma), 100 units/mL of penicillin, and 100 μg/mL of streptomycin (Invitrogen). HA-tagged NPHP1 expression vectors were transfected to mIMCD-3 cells using FuGENE HD (Promega) following the manufacturer’s instruction. Immunostaining and imaging procedures are described in [37].

### Electroretinogram (ERG)

Mice were dark-adapted overnight before ERG recording. Under dim red light illumination, mice were anesthetized by intraperitoneal injection of a ketamine/xylazine mixture (87.5 mg/kg and 12.5 mg/kg, respectively), and pupils were dilated with 1% tropicamide ophthalmic solution (Akorn) for 2-3 minutes. Mice were placed on a Celeris D430 rodent ERG testing system (Diagnosys) with its heater on to maintain animals’ body temperature. After applying GenTeal Tears Lubricant Eye Gel (Alcon), light-guide electrodes were positioned on both eyes. For dim light scotopic ERG, responses were measured with 15 flashes of 0.01 cd.sec/m^2^ stimulating light (color temperature: 6500K). Dark-adapted Standard Combined Response (SCR) was measured with 15 flashes of 3.0 cd.sec/m^2^ stimulating light (color temperature: 6500K). For photopic ERG, mice were light-adapted for 10 minutes (background light; 9.0 cd.sec/m^2^) and responses were measured with 15 flashes of 3.0 cd.sec/m^2^ white light (6500K). After recording, animals were allowed to recover on a heating pad.

### RNA extraction and reverse transcription (RT)-PCR

Tissue collection and RNA extraction procedures were described previously [25]. *Nphp1* cDNA fragments between exons 2 and 11 were PCR-amplified using Universe High-Fidelity Hot Start DNA polymerase (Bimake) and two primers (F1-mNPHP1-RT and R1-mNPHP1-RT) listed in **Table 1**. PCR products were directly sequenced by Sanger sequencing using the F1-mNPHP1-RT primer. Band intensities were measured using the Image Lab software (Bio-Rad).

### Immunoprecipitation

HEK 293T/17 cells were obtained from ATCC (#CRL-11268) and cultured in Dulbecco’s Modified Eagle’s Medium (DMEM; Invitrogen) supplemented with 10% fetal bovine serum (Sigma), 100 units/mL of penicillin, and 100 μg/mL of streptomycin (Invitrogen). GFP-NPHP2 and GFP-NPHP5 expression vectors were transiently transfected to 293T/17 cells together with pCS2FLAG empty vectors (as a control) or pCS2FLAG-NPHP1 variants using FuGENE HD (Promega), following the manufacturer’s instruction. Protein extracts were prepared and subjected to immunoprecipitation with anti-FLAG M2 magnetic beads (Sigma) as previously described [25]. SDS-PAGE and immunoblotting were performed as described above.

## Results

### Mislocalization of inner segment plasma membrane-associated proteins in *Nphp1*^*gt/gt*^ retinas

To test the requirement of NPHP1 for compartmentalized protein localization in photoreceptors, we probed the localization of various IS and OS resident proteins in *Nphp1*^*gt/gt*^ mouse retinas. Our prior study showed that IS plasma membrane proteins STX3, STXBP1, SNAP25, and IMPG2 were the most susceptible to mislocalization in CEP290-deficient photoreceptors [25]. Therefore, we focused our efforts on these proteins. In normal photoreceptors, these proteins were strictly restricted to the IS (hereafter our use of the term IS encompasses all parts of photoreceptors except the OS) (**Fig 1**). In 18-day old *Nphp1*^*gt/gt*^ mutants, however, all of these proteins showed significant mislocalization to the OS (**Fig 1A-D**; see **Fig 1E** for quantification). STXBP1 showed the most severe mislocalization with approximately 1/3 of proteins mislocalized to the OS. Consistent with the previous finding [22], mislocalization of RHO to the IS was relatively mild in *Nphp1*^*gt/gt*^ mutants and no mislocalization was detected with other OS resident proteins examined (PRPH2, ROM1, ABCA4, and PDE6B) (**Fig 1 and S3 Fig**).

**Fig 1.**
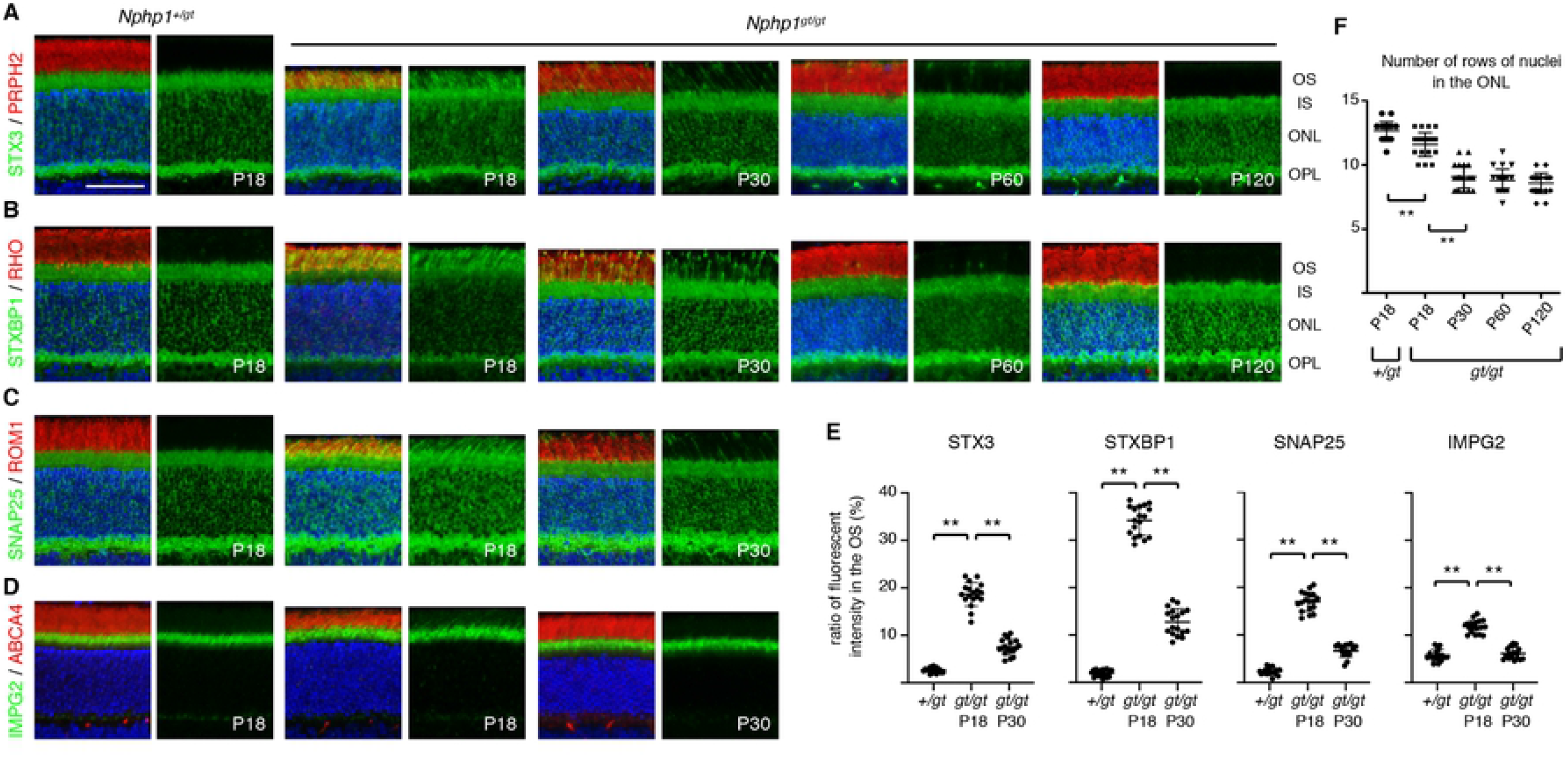
Temporary impairment of protein confinement and retinal degeneration in *Nphp1*^*gt/gt*^ mice. (A-D) Transient mislocalization of STX3 (A), STXBP1 (B), SNAP25 (C), and IMPG2 (D) in *Nphp1*^*gt/gt*^ retinas. Retinal sections from *Nphp1*^*+/gt*^ and *Nphp1*^*gt/gt*^ mice were stained with STX3, STXBP1, SNAP25, and IMPG2 antibodies (green). PRPH2, RHO, ROM1, and ABCA4 (red) were labeled as a marker of the OS. Merged images are shown on the left. DAPI (blue) was used to counterstain the nuclei. Scale bar denotes 50 μm. IS: inner segment, ONL: outer nuclear layer, OPL: outer plexiform layer. OS: outer segment. (E) Quantification of IS membrane protein mislocalization to the OS. Depicted are the ratios of integrated fluorescence intensities in the OS relative to the photoreceptor cell layer. Mean ± standard deviation (SD; error bars) is marked by horizontal lines (n=3 mice; 2 sections/mouse and 3 areas/section). Asterisks indicate statistical significance (one-way ANOVA followed by Tukey’s multiple comparison test; *p* < 0.01). (F) Temporary retinal degeneration in *Nphp1*^*gt/gt*^ retinas. The number of rows of photoreceptor cell nuclei was counted in *Nphp1*^*+/gt*^ and *Nphp1*^*gt/gt*^ mice at the central retina (n=4 mice; 3 sections/mouse; 2 locations/section). Mean ± SD is shown. Asterisks indicate statistical significance (one-way ANOVA; *p* < 0.01).

We then examined whether IS membrane proteins progressively accumulated in the OS as in *Cep290* mutant retinas [25]. To our surprise, localization of all proteins examined (STX3, STXBP1, SNAP25, and IMPG2) was significantly improved in 30-day old *Nphp1*^*gt/gt*^ mutants and only a subset of photoreceptors displayed mislocalization (P30 in **Fig 1A-E**). Localization of these proteins was even further improved by P60 and became indistinguishable from normal controls by P120. Mislocalization of RHO, PRPH2, ROM1, ABCA4, and PDE6B was not detected after P30 (**Fig 1 and S3 Fig**).

To test whether loss of NPHP1 more preferentially affects cone OS proteins, we examined the localization of OPN1MW (cone opsin) and GNAT2 (cone transducin α) in *Nphp1*^*gt/gt*^ mutants (**Fig 2**). In 18-day old *Nphp1*^*gt/gt*^ mice, cone OSs appeared to be short and disorganized. OPN1MW mostly localized to the OS but a small proportion was found mislocalized to the IS (**Fig 2A and C**). In contrast, localization of GNAT2 was not altered in *Nphp1*^*gt/gt*^ retinas (**Fig 2B and C**). At P30 and P120, cone OSs appeared to be normal in length and shape, and no mislocalization of cone OS proteins was detected.

**Fig 2.**
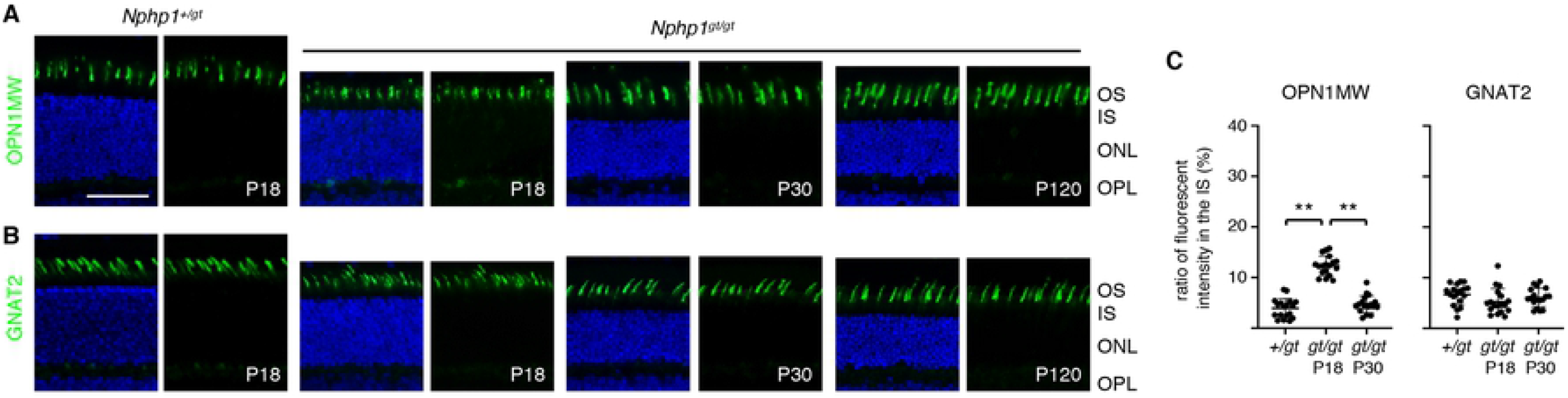
Localization of cone OS proteins in *Nphp1*^*gt/gt*^ retinas. (A-B) Cone OS proteins, OPN1MW (A) and GNAT2 (B), were immunostained in *Nphp1*^*+/gt*^ and *Nphp1*^*gt/gt*^ retinal sections. Mild mislocalization of OPN1MW was observed at P18 but not at P30 in *Nphp1*^*gt/gt*^ retinas. Scale bar denotes 50 μm. (C) Quantification of cone OS protein mislocalization to the IS. Others are the same as in Fig 1E.

### Retinal degeneration in *Nphp1*^*gt/gt*^ mice

We examined photoreceptor degeneration in *Nphp1*^*gt/gt*^ mutants. To this end, we counted the number of rows of photoreceptor cell nuclei at the central retina (300-600 μm from the optic nerve head). Consistent with the previous finding [22], there was a slight loss (∼1 row) of photoreceptor cells by P18 (mean ± SD: 12.7 ± 0.7 in *Nphp1*^*+/gt*^ vs. 11.6 ± 0.9 in *Nphp1*^*gt/gt*^; *p*<0.01; n=4) (**Fig 1F**). Thinning of the outer nuclear layer became more obvious and 3-4 rows of photoreceptors were lost by P30 (mean ± SD: 9.1 ± 0.9 at P30; *p*<0.01 compared with *Nphp1*^*gt/gt*^ at P18; n=4). However, retinal degeneration stopped thereafter and ∼70% of photoreceptors were retained until 4 months of age (the last time point examined) in *Nphp1*^*gt/gt*^ mice (8.8 ± 0.9 at P60 and 8.6 ± 0.8 at P120; n=4) (**Fig 1F**).

### Normal retinal functions in *Nphp1*^*gt/gt*^ mice

We next examined whether *Nphp1*^*gt/gt*^ mouse retinas were functionally normal despite the initial protein mislocalization and partial loss of photoreceptors (**Fig 3**). In both scotopic and photopic conditions, electroretinogram (ERG) of 2-month old *Nphp1*^*gt/gt*^ mice was comparable to that of control animals (n=4). These data indicate that both rods and cones recover from the initial protein confinement defect and become fully functional in *Nphp1*^*gt/gt*^ mice.

**Fig 3.**
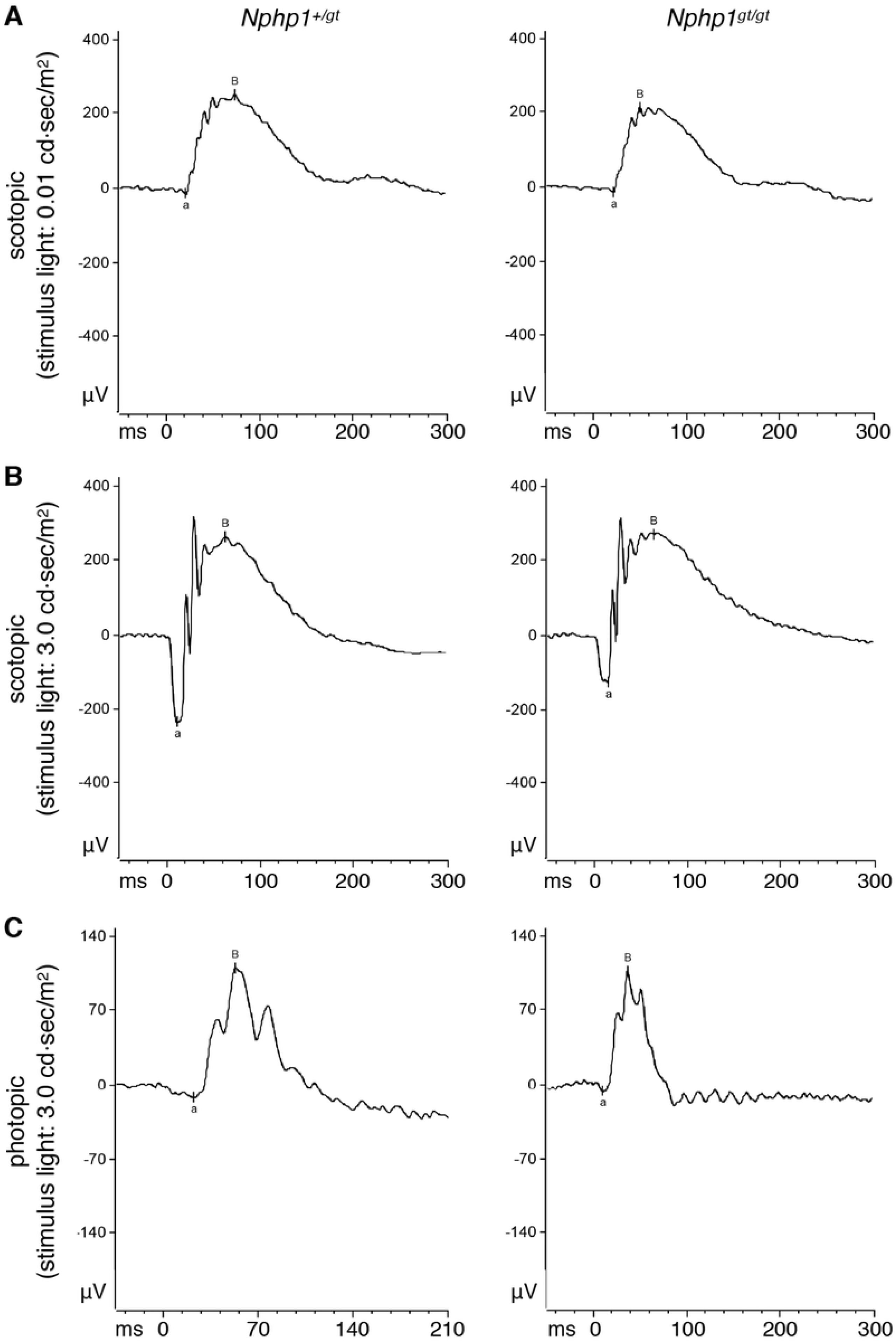
*Nphp1*^*gt/gt*^ mouse retinas respond normally to light. Scotopic (A and B) and photopic (C) ERG recordings of 2-month old *Nphp1*^*+/gt*^ and *Nphp1*^*gt/gt*^ mice are shown. Stimulus light intensities are shown on the left.

### Production of functional mRNAs from the *Nphp1*^*gt*^ allele

Spontaneous amelioration of protein mislocalization and temporary retinal degeneration in *Nphp1*^*gt/gt*^ mice sharply contrast with the severe retinal degeneration phenotype observed in *Nphp1*^*del20/del20*^ mice [5]. To gain insights into the molecular basis of these striking differences, we examined possible expression of normal or mutant variants of NPHP1 in *Nphp1*^*gt/gt*^ mice. More specifically, we directed our attention to non-canonical mRNA splicing events that could result in the production of functional proteins. Basal exon skipping and nonsense-associated altered splicing are mechanisms, through which cells eliminate nonsense or frameshift mutation-containing exons during the pre-mRNA splicing. The resulting mRNAs maintain the reading frame of the gene and produce near-full-length proteins [45-50]. Neighboring exon(s) may be skipped as well if omitting the mutated exon alone does not prevent frameshift. These non-canonical splicing events have been reported in the human *CEP290* gene and account for the unexpectedly mild phenotypes observed in certain *CEP290*-LCA patients [47-50]. We recently showed that nonsense-associated altered splicing also occurs in a mouse model *CEP290*-LCA [25]. The *Nphp1*^*gt*^ allele was generated by an insertion of a gene-trap into exon 4 (**S1 Fig**) [22], and we examined whether altered splicing occurred in *Nphp1*^*gt/gt*^ mice.

To assess exon skipping, RNAs were extracted from 1.5-month old *Nphp1*^*+/+*^ and *Nphp1*^*gt/gt*^ mouse eyes, and cDNA fragments were PCR amplified using primers specific to *Nphp1* exons 2 and 11 (forward and reverse primers, respectively). In *Nphp1*^*+/+*^ mice, a single PCR product containing all exons between exons 2 and 11 was amplified (991 bp; based on the transcript ID ENSMUST00000028857.13) (**Fig 4A**; left), suggesting no basal exon skipping. A slightly smaller fragment (805 bp) was amplified from *Nphp1*^*gt/gt*^ mice. Sequence analyses of these PCR products revealed that the 805-bp fragment contained *Nphp1* coding sequences but lacked exons 3 and 4 as well as the gene-trap (**Fig 4B**). These data indicate that the entire gene-trap plus *Nphp1* exons 3 and 4 were skipped during pre-mRNA splicing. *Nphp1* exons 3 and 4 are 61 bp and 125 bp long, respectively. Skipping of these two exons results in an in-frame deletion of 62 amino acids (aa; from Cys49 to Lys110 (p.C49_K110del)) within the N-terminal coiled-coil domain (**Fig 4C**). These data demonstrate that nonsense-associated altered splicing does occur in *Nphp1*^*gt*^ transcripts.

**Fig 4.**
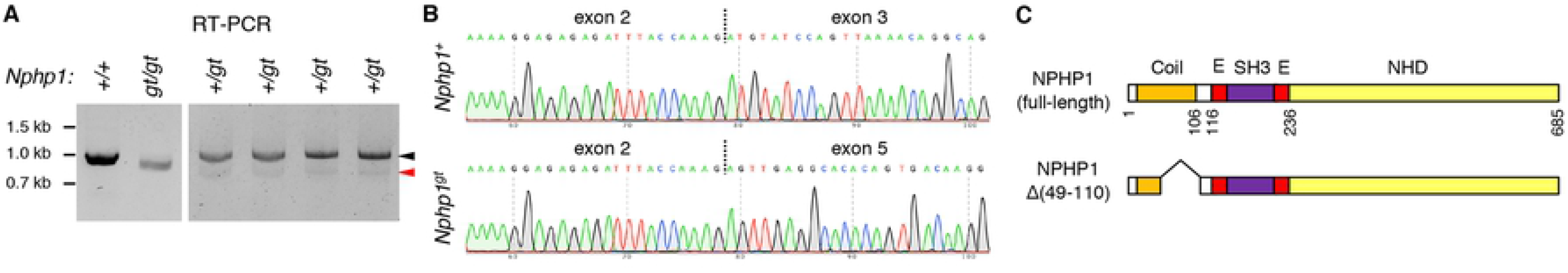
A small quantity of functional mRNA is produced from the *Nphp1*^*gt*^ allele via nonsense-associated altered splicing. *(A) Nphp1* RT-PCR data from *Nphp1*^*+/+*^, *Nphp1*^*gt/gt*^, and *Nphp1*^*+/gt*^ retinas. Black and red arrowheads indicate PCR products from the wild-type and the *Nphp1*^*gt*^ allele, respectively. Each animal’s *Nphp1* genotype is shown at the top. Locations of DNA size markers are shown on the left. (B) Chromatogram of RT-PCR product sequencing reactions around exon-exon junctions. Vertical dotted lines indicate exon-exon junctions. (C) Schematic representation of NPHP1 protein products encoded by wild-type and *Nphp1*^*gt*^ alleles. Major structural domains are depicted (Coil: coiled-coil domain; E: Glu-rich domain; SH3: SH3 domain; NHD: nephrocystin homology domain). Amino acid positions are shown below full-length NPHP1 and based on the translated amino acid sequence of BC118953.

We noticed that the band intensity of the 805-bp fragment was significantly lower than that of wild-type fragments (**Fig 4A**; left). To better assess the quantity of *Nphp1*^*gt*^ mRNAs produced by nonsense-associated altered splicing, we performed PCR using cDNAs from *Nphp1*^*+/gt*^ heterozygous mice and stopped PCR amplification after 30 cycles (i.e., before saturation) (**Fig 4A**; right). Using wild-type bands as an internal control, densitometric analysis of the PCR products indicated that the quantity of *Nphp1*^*gt*^ altered splicing products was less than 10% of wild-type mRNAs (mean ± SD: 6.3 ± 1.0%; n=4). To test whether the nonsense-associated altered splicing is an age-dependent phenomenon and underlies the temporary retinal phenotypes, RNAs were extracted from 10-day old *Nphp1*^*+/+*^, *Nphp1*^*+/gt*^, and *Nphp1*^*gt/gt*^ mouse eyes, and RT-PCR was conducted using the same PCR primers. RT-PCR results from young animals were similar to those of adult mice (**S4 Fig**), indicating that nonsense-associated altered splicing occurs similarly in both developing and matured retinas.

### Expression and localization of the NPHP1 deletion mutant encoded by the *Nphp1*^*gt*^ allele

We characterized the NPHP1 mutant protein encoded by the *Nphp1*^*gt*^ allele (hereafter NPHP1 Δ(49-110)). With the antibodies available to us, we were not able to detect endogenous NPHP1 convincingly in the retina by immunoblotting. Therefore, we used protein extracts from testes, in which ciliary proteins were enriched (**Fig 5A**). While full-length NPHP1 with a calculated molecular weight of 77 kDa was readily detected in *Nphp1*^*+/+*^ testes (red arrowhead), we were not able to detect any unique protein band in *Nphp1*^*gt/gt*^ testes.

**Fig 5.**
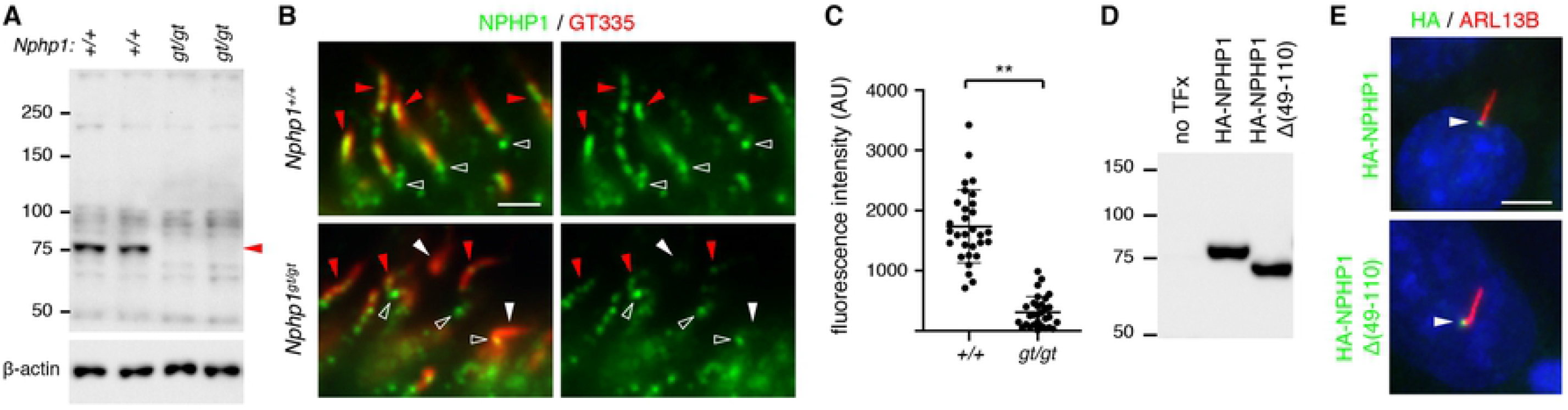
Expression and localization of NPHP1 Δ(49-110) deletion mutants. (A) Immunoblotting of NPHP1 (red arrowhead) in *Nphp1*^*+/+*^ and *Nphp1*^*gt/gt*^ mouse testes. Fifty μg of proteins were loaded per lane. β-actin was used as a loading control. (B) Localization of NPHP1 Δ(49-110) to the connecting cilium. Connecting cilia are marked by GT335 polyglutamylation antibodies (red). NPHP1 immunoreactivity (green) is detected at the connecting cilium (red arrowheads) in both *Nphp1*^*+/+*^ and *Nphp1*^*gt/gt*^ retinas but significantly reduced in *Nphp1*^*gt/gt*^ retinas. White arrowheads indicate connecting cilia with no NPHP1 immunoreactivity. Open arrowheads denote cross-reacting protein(s) around the basal body. Scale bar represents 1 μm. (C) Quantification of NPHP1 immunofluorescence intensities (in arbitrary units; AU) at the connecting cilium. Mean ± SD is marked by horizontal lines (n=30; 2 mice). Asterisks indicate statistical significance (*t*-test; *p* < 0.001). (D) Expression of HA-tagged NPHP1 and NPHP1 Δ(49-110) in transiently transfected 293T/17 cells. (E) Localization of NPHP1 Δ(49-110) to the transition zone in mIMCD-3 cells. HA-NPHP1 and HA-NPHP1 Δ(49-110) were transiently transfected to mIMCD-3 cells and their localization was probed with anti-HA antibodies (green; white arrowhead). ARL13B (red) was labeled as a marker of primary cilia. Scale bar represents 5 μm.

We then examined the localization of NPHP1 in the retina by immunohistochemistry (**Fig 5B**). As previously reported [4, 5], NPHP1 was found at the connecting cilium in normal photoreceptor cells (red arrowhead). Signals around the basal body (open arrowhead) were from cross-reacting protein(s). Consistent with the immunoblotting data, NPHP1 immunoreactivity at the connecting cilium was drastically reduced in *Nphp1*^*gt/gt*^ retinas (**Fig 5B and C**). However, low-level immunoreactivity was detected in a subset of cells (**Fig 5B**; red arrowhead), suggesting that NPHP1 Δ(49-110) mutants are expressed and localize to the connecting cilium.

To confirm the localization of NPHP1 Δ(49-110) to the connecting cilium, we generated expression plasmids encoding the full-length and the deletion mutant form of NPHP1 with an N-terminal HA tag and examined their localization in ciliated mIMCD-3 cells. As shown in **Fig 5D and E**, HA-NPHP1 Δ(49-110) was as efficiently produced as HA-NPHP1 and localized to the transition zone in mIMCD-3 cells. These data indicate that the mutant protein retains its ability to localize to the connecting cilium.

### NPHP1 Δ(49-110) mutants retain the ability to interact with other NPHP proteins

We then examined whether the 62-aa deletion had any impact on protein-protein interactions of NPHP1. NPHP1 is previously shown to interact with AHI1, NPHP2 (also known as INVS), NPHP4, NPHP5 (IQCB1), and NPHP8 (RPGRIP1L) [10, 51-54]. Of these, NPHP1 directly binds to AHI1 and NPHP4 via its SH3 domain and the C-terminal 131 residues, respectively [10, 51, 53, 54]. Since these regions are preserved in the NPHP1 Δ(49-110) mutant and sufficient to mediate NPHP1 interactions with AHI1 and NPHP4, we focused our efforts on interactions with NPHP2, NPHP5, and NPHP8. To this end, we transiently transfected FLAG-tagged NPHP1 variants (full-length NPHP1, NPHP1 Δ(49-110), and NPHP1 aa242-685) with GFP-tagged NPHP2 and NPHP5, and performed immunoprecipitation (IP) using anti-FLAG antibodies (**Fig 6**). For NPHP8, endogenous proteins were probed. As shown in **Fig 6**, both NPHP1 Δ(49-110) and NPHP1 aa242-685 were able to pull down all three NPHP proteins, indicating that the C-terminal nephrocystin homology domain (NHD) alone was sufficient to interact with these proteins either directly or indirectly. These data show that the NPHP1 Δ(49-110) mutant retains most, if not all, of its protein-protein interaction capabilities.

**Fig 6.**
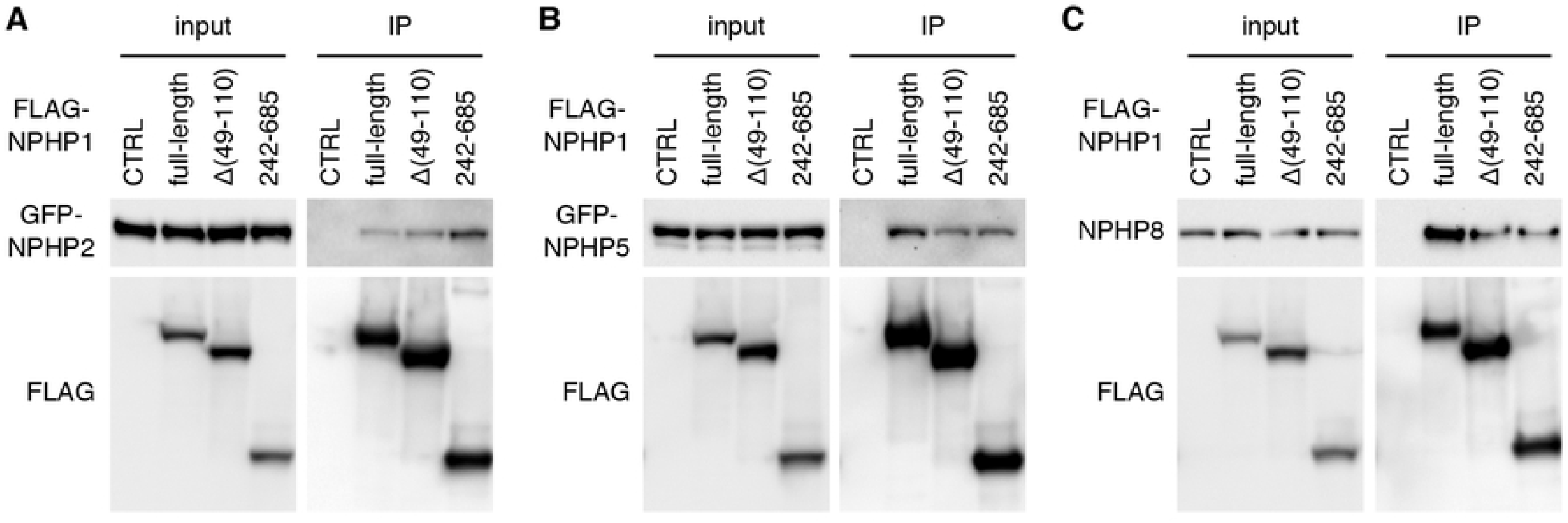
Interaction of NPHP1 Δ(49-110) with other NPHP proteins. Expression vectors of FLAG-tagged NPHP1 (full-length), NPHP1 Δ(49-110), and NPHP1 aa242-685 were co-transfected with either pEGFP-NPHP2 (A) or pEGFP-NPHP5 (B) to 293T/17 cells, and protein extracts were subjected to immunoprecipitation (IP) with anti-FLAG antibodies. Empty vectors were used as a negative control (CTRL). For NPHP8 (C), endogenous NPHP8 was probed.

### *Cep290* genetically interacts with *Nphp1* and modifies retinal phenotypes in *Nphp1*^*gt/gt*^ mice

Finally, we tested whether additional mutations in other ciliary gate-related genes could modify the severity of retinal degeneration in *Nphp1*^*gt/gt*^ mice. To this end, we crossed *Nphp1*^*gt/gt*^ mice to *Cep290*^*fl/fl*^;*iCre75* mice [25, 30, 31] and compared protein mislocalization and retinal degeneration phenotypes in *Nphp1*^*gt/gt*^;*Cep290*^*+/+*^;*iCre75* and *Nphp1*^*gt/gt*^;*Cep290*^*+/fl*^;*iCre75* animals: all animals analyzed in this study were *iCre75* hemizygotes and *iCre75* genotype is omitted for brevity. At 1 month of age, mislocalization of IS plasma membrane proteins STX3, STXBP1, and SNAP25 was significantly increased in *Nphp1*^*gt/gt*^;*Cep290*^*+/fl*^ mice compared with that in *Nphp1*^*gt/gt*^;*Cep290*^*+/+*^ mice (**Fig 7**). RHO mislocalization was also detected in *Nphp1*^*gt/gt*^;*Cep290*^*+/fl*^ mice, but it was relatively mild compared to that of IS membrane proteins. Consistent with the increased protein mislocalization, reduction of the outer nuclear layer was evident in *Nphp1*^*gt/gt*^;*Cep290*^*+/fl*^ mice. At 2 months of age, when *Nphp1*^*gt/gt*^;*Cep290*^*+/+*^ mice exhibited no mislocalization and no degeneration, *Nphp1*^*gt/gt*^;*Cep290*^*+/fl*^ mice exhibited severe retinal degeneration with only 3-5 rows of photoreceptor cell nuclei remaining. In *Nphp1*^*gt/gt*^;*Cep290*^*fl/fl*^ double mutants, more than 90% of photoreceptors were lost by P30, which is faster than what we observed in *Cep290*^*fl/fl*^ single mutants using the same *Cre* driver [25]. These data indicate that *Nphp1* and *Cep290* genetically interact and that reduction of *Cep290* gene dose exacerbates protein confinement defects and causes continuous retinal degeneration in *Nphp1*^*gt/gt*^ mice.

**Fig 7.**
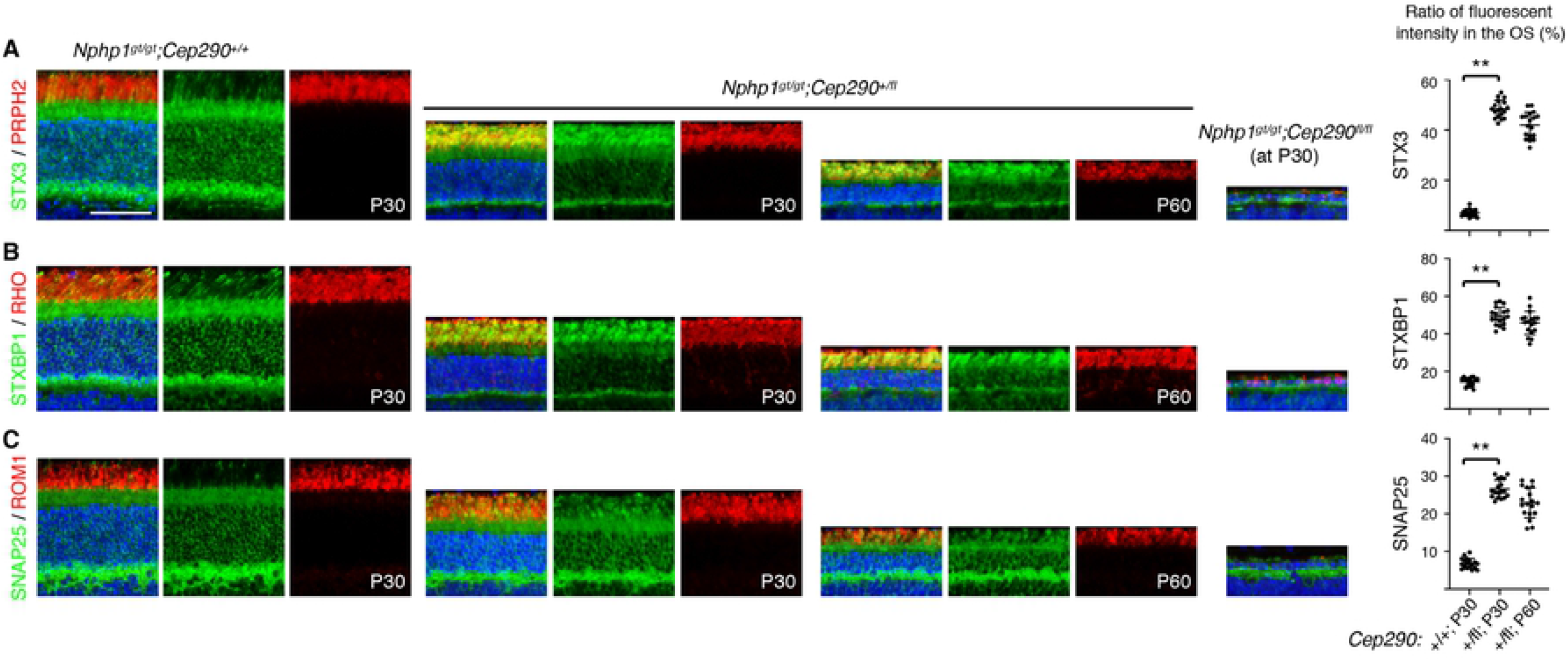
Genetic interaction of *Nphp1* with *Cep290*. Mislocalization of STX3 (A), STXBP1 (B), and SNAP25 (C) in *Nphp*^*gt/gt*^*;Cep290*^*+/+*^, *Nphp*^*gt/gt*^*;Cep290*^*+/fl*^, and *Nphp*^*gt/gt*^*;Cep290*^*fl/fl*^ retinas (green). PRPH2, RHO, and ROM1 (red) were labeled to mark the OS. The ratio of mislocalized proteins was quantified as in Fig 1 and depicted on the right. Mean ± SD is marked by horizontal lines (n=3 mice; 2 sections/mouse and 3 areas/section). Asterisks indicate statistical significance (one-way ANOVA followed by Tukey’s multiple comparison test; *p* < 0.01).

## Discussion

In this work, we characterize retinal degeneration in a mouse model of NPHP1 and demonstrate the requirement of NPHP1 for compartmentalized protein localization in photoreceptors. The mutant allele in the *Nphp1*^*gt*^ model is a hypomorph rather than a null because of the production of a small number of functional mRNAs derived from nonsense-associated altered splicing. The resulting mRNAs encode a near-full-length protein with a 62-aa in-frame deletion. The mutant protein appears to retain most, if not all, functions of the full-length protein, as it correctly localizes to the ciliary transition zone and interacts with its known binding partners. However, the quantity of the functional mRNAs produced by altered splicing is very low (less than 10% of wild type), and the protein products are undetectable or barely detectable in the testis and the retina with the reagents available to us. Although we were not able to detect the mutant protein in the testis, it is noteworthy that all *Nphp1*^*gt/gt*^ males (n=12) used for breeding produced offspring with normal litter size (5-12 pups/litter). This is contrary to the sterility of *Nphp1*^*del20/del20*^ males [21]. Therefore, our data suggest that the *Nphp1*^*gt*^ allele is a hypomorph and that the phenotypic differences between the two NPHP1 mouse models are due to a difference in the residual activities of the two mutant alleles rather than genetic backgrounds.

Our study shows that NPHP1 is involved in membrane protein confinement at the connecting cilium. In developing *Nphp1*^*gt/gt*^ retinas, confinement of IS plasma membrane proteins is compromised, leading to an infiltration of IS membrane proteins into the OS. Among the OS proteins examined, only opsins (RHO and OPN1MW) showed mislocalization to the IS and the degree of mislocalization was relatively mild compared to that of IS plasma membrane proteins. This phenotype is very similar to what is observed in *Cep290*^*fl*^ mice [25]. It also should be noted that, despite more severe RHO mislocalization and faster degeneration, IS membrane protein mislocalization does not occur in *Ift88* mutants (**S5 Fig**). This suggests that IS membrane protein mislocalization is not a consequence of the degenerative process occurring in dying cells. Therefore, although it has been examined in only two disease models thus far, our findings are consistent with the idea that accumulation of IS membrane proteins in the OS is a common pathomechanism of retinal degenerations associated with ciliary gate defects [3]. Further investigation is needed to validate this idea in other ciliary gate-related retinal degenerations. It is also noteworthy that although Bardet-Biedl syndrome (BBS) proteins are not part of the ciliary gate complex, retinal degeneration in BBS shares the same disease mechanism [38, 55, 56]. Future studies should be directed to how this molecular phenotype leads to the death of photoreceptor cells.

Interestingly, however, the requirement of NPHP1 in photoreceptors is developmental stage dependent. In mice, the connecting cilium assembles at P3-5 and the OS develops afterward until P18-20 [57, 58]. Protein mislocalization occurs in *Nphp1*^*gt/gt*^ retinas when OSs are actively elongating, but normal distribution patterns are restored after P18. Some photoreceptors die during this period, but retinal degeneration stops as the protein confinement defect is ameliorated. Retinal functions and anatomy are normal in adult *Nphp1*^*gt/gt*^ mice except ∼30% thinner ONL due to the earlier loss of photoreceptors. RT-PCR data indicate that nonsense-associated altered splicing of *Nphp1*^*gt*^ pre-mRNAs occurs in not only fully matured but also developing retinas. Therefore, age-dependent differences in altered splicing are not likely a contributing factor. One speculation is that, although ciliary gate functions are essential throughout the life of photoreceptors, more vigorous NPHP1 activities might be needed when OSs are rapidly elongating compared to when the OS elongation is completed.

Our study suggests that additional mutations in other ciliary gate-related genes may affect the penetrance of retinopathy in human NPHP1 patients. As mentioned in the Introduction, only 6-10% of NPHP1 patients manifest retinal anomalies [19, 20, 59]. The most frequent mutations in human *NPHP1*, found in 65-80% of NPHP1 patients, are large (∼250 kb) homozygous deletions that eliminate the majority of *NPHP1* exons [16-18, 60]. Since these mutations are expected to be null, NPHP1 does not appear to be essential for photoreceptor health in humans. A previous study showed that removal of one allele of *Ahi1*, which encodes a component of the ciliary gate complex [10], significantly increases RHO mislocalization and accelerates retinal degeneration in *Nphp1*^*gt/gt*^ mice [22]. In the present work, we show that reducing the gene dose of *Cep290* in *Nphp1*^*gt/gt*^ mice exacerbates protein mislocalization and causes retinal degeneration that continues in mature retinas. These findings suggest that *NPHP1* genetically interacts with other ciliary gate-related genes and that the manifestation of retinopathy in NPHP1 patients may be affected by the presence of additional mutations in those genes.

Our study also suggests that mutation-induced non-canonical splicing might be more common than what has been appreciated. As mentioned above, some *CEP290*-LCA patients exhibit relatively mild phenotypes despite the presence of truncation mutations that should eliminate a large proportion of the protein [47-50]. The unexpectedly mild phenotypes are due to the production of near-full-length proteins derived from non-canonical mRNA splicing that skips exons with truncating mutations. One of the interesting features of inherited retinal degenerations is the wide spectrum of phenotypic severity despite mutations in the same gene [20, 61-67]. Although some of these variations can be explained by disease-causing mutations *per se* (e.g. missense mutations that partly reduce protein activities) and genetic modifiers, nonsense-associated altered splicing may contribute to the variation and underlie the mild end of the phenotypic spectrum of disease.

Finally, our study urges a more thorough investigation of mRNAs produced in mutant animals. When loss-of-function animal models are generated, inducing frameshift is a frequently used strategy. For example, CRISPR-based gene knockout approaches utilize the tendency of introducing indels at double-strand breaks, which results in a frameshift. In more conventional recombination-based approaches, exons, deletion of which causes frameshift, are preferentially selected when designing targeting vectors. Accumulating evidence indicates that exons containing premature termination codons are often skipped as a compensatory mechanism to cope with induced mutations [68-70]. Our results show that gene-traps can be skipped as well during splicing. Therefore, a thorough investigation of mRNAs produced from mutant alleles is warranted before beginning extensive studies and making conclusions without knowing the precise consequences of the mutation.

## Conclusions

Our study shows that NPHP1 is required to maintain compartmentalized protein localization during the photoreceptor terminal differentiation, particularly to prevent infiltration of IS plasma membrane proteins into the OS. However, this requirement is developmental stage-dependent, and NPHP1 becomes less crucial as the OS elongation is completed. Retinal degeneration occurs when NPHP1 deficiency causes protein mislocalization but stops as normal localization patterns are restored. These findings suggest that accumulation of IS membrane proteins in the OS is part of the disease mechanisms underlying the *NPHP1*-associated retinopathy. Our study further shows that reduction of *Cep290* gene dose exacerbates protein confinement defects and retinal degeneration in *Nphp1* mutants. These findings suggest that additional mutations in other ciliary gate-related genes may influence the penetrance of retinopathy in human NPHP1 patients.

## Acknowledgments

We thank Drs. Val C. Sheffield and William Y. Tsang for the gifts of pEGFP-NPHP2 and pEGFP-NPHP5. This work was supported by National Institutes of Health grants R01-EY022616 and R21-EY027431 (to S.S.), National Institutes of Health Center Support grant P30-EY025580 (to the University of Iowa), and Research to Prevent Blindness Unrestricted Grant (to the Department of Ophthalmology and Visual Sciences, University of Iowa).

## Abbreviations

DAPI: 4′,6-Diamidine-2′-phenylindole dihydrochloride
DMEM: Dulbecco’s modified Eagle’s medium
CRISPR: clustered regularly interspaced short palindromic repeats
ERG: electroretinogram
IS: inner segment
JBTS: Joubert syndrome
LCA: Leber congenital amaurosis
MKS: Meckel-Gruber syndrome
NPHP: nephronophthisis
ONL: outer nuclear layer
OPL: outer plexiform layer
OS: outer segment
RT-PCR: reverse transcription polymerase chain reaction
SD: standard deviation

